# Moving beyond mean thermal death times to assess organismal responses to stressful temperatures

**DOI:** 10.64898/2026.01.15.699638

**Authors:** Garrison W. Bullard, Lauren B. Buckley, Joel G. Kingsolver

## Abstract

Predicting survival of ectotherms in stressful and variable thermal environments is an essential challenge in this era of heat waves and climate change. Recent thermal death time (TDT) models, based on an exponential relationship between average time to death (or failure) t_f_ and temperature, enable accounting for average survival responses to both the magnitude and duration of stressful temperatures. However, extending these deterministic and probabilistic models to predict patterns of survival in fluctuating temperatures currently requires additional assumptions: e.g. that injury accumulation due to heat stress is additive across temperatures, and that the shape of the cumulative survival curve does not change with temperature. We evaluate these assumptions and their consequences by using a parametric survival model and available data on failure (knockdown) times of adult *Drosophila.* We find that the variance in log(t_f_) increases with increasing constant temperatures in most *Drosophila* species, resulting in changes in shape of the failure density and survival curves across temperatures. We compare predictions of three deterministic and probabilistic models that differ in their TDT assumptions using *D. melanogaster* data in fluctuating (but stressful) temperatures. All three models consistently underestimate observed median failure times except at extremely high temperatures, suggesting non-additivity of heat injury accumulation. Our parametric model, incorporating temperature-dependent variance, provides more accurate predictions of cumulative survival curves in fluctuating temperatures. Our findings highlight the importance of understanding both mean and variation in failure times, and how these change across temperatures, for modeling survival in fluctuating thermal environments.

## INTRODUCTION

Increases in environmental variability, heat waves, and other extreme thermal events due to climate change are amplifying the need to understand organismal responses to high and fluctuating temperatures (1–3). Integrating widely available metrics of organismal thermal tolerance (4) with environmental data provides a convenient way to estimate the thermal risk posed by climate change (5,6). Tolerance has been quantified using both static (time to loss of function at a constant stressful temperature) and dynamic (temperature at loss of function as temperature is ramped) approaches, but metrics of thermal tolerance using either approach are highly sensitive to methodology (7–9). Beyond methodological challenges, the probability of survival depends on both the intensity and duration of thermal stress (8,10). Thermal sensitivity also depends on the thermal history the organism has experienced, with past stressful temperatures alternately allowing for acclimation or the accumulation of stress and cool temperatures allowing for recovery (11,12).

Efforts to account for both temperature and exposure time have led to a shift toward considering a two-dimensional ‘tolerance landscape’ rather than a single thermal tolerance metric (10). One valuable approach to predicting survival in fluctuating temperatures uses thermal death times (TDTs): the time to death (or failure) as a function of temperature (1,3). Data from a variety of study systems indicate that mean time to death declines exponentially with increasing temperature (13–15). This yields a linear relationship between temperature T and log of time to death or failure t_f_: log(t_f_) ∼ bT, where b is the slope. Empirical studies have defined failure in terms of death, knockdown, coma or other events; for simplicity, here we will refer to all of these in terms of failure and failure time t_f_, and survival as the absence of failure. Recent studies have used a variety of statistical approaches to estimate the relationship between log(t_f_) and temperature for different constant temperatures (16–20). However, applying these models based on constant temperature data to the case of fluctuating temperatures requires additional information or assumptions about how time and temperature jointly determine survival (see below).

The basic TDT framework has motivated the development of both deterministic and stochastic (probabilistic) models of survival in changing temperatures. Deterministic models of failure in stressful temperatures assume that failure of an organism occurs when accumulation of injury or damage exceeds a critical threshold. For example, in the model proposed by Jørgensen et al (1), the rate of injury depends on temperature, and the injury accumulation is assumed to be additive across temperatures (1,9). Several experimental studies of knockdown times in *Drosophila* support this assumption, but the range of temperature conditions in which this applies is unknown (1,9). Deterministic models can be used to predict average failure times in fluctuating temperatures when failure is due to stress; however these models do not consider the stochastic nature of failure, so cannot predict survival probabilities or variation in failure times.

Conversely, probabilistic models explicitly consider the cumulative survival curve S(t) and how it varies with temperature T and time t. Applying such models to fluctuating temperatures requires assumptions about S(t) and the distribution of failure times. One straightforward approach is to assume that the failure rate at time t depends only on current temperature T(t): in this case failure time is exponentially distributed with a survival rate λ = 1/t_f_, such that failure times and survival curves can be computed directly from the TDT relationship. This method may apply to immature life stages at non-stressful temperatures, but in general survival will depend on age, past thermal history (including thermal stress) and other factors (10,21,22). The influential TDT model of Rezende et al (3) does not assume a specific parametric form for the cumulative survival curve S(t), but assumes that there is an ‘average’ S(t) that applies across all temperatures of interest (3). However, the shape of S(t)-- and variation in failure times– is known to be influenced by both biological mechanisms and external environmental factors (22–24).

These deterministic and probabilistic models provide related but distinct methods for using data on death or failure times over a range of constant, stressful temperatures to predict survival in fluctuating temperatures, and have been applied to *Drosophila* (1,3) and other ectotherms (25–28). However, whether and how cumulative survival curves change across stressful temperatures has not been examined. As we will show, this has important consequences for predicting survival probabilities in fluctuating conditions. In addition, the accuracy of predictions from current deterministic vs probabilistic models has not been directly compared.

Here, we explore these issues and their consequences for predicting survival in constant and fluctuating temperature conditions. First, we use a simple probability model to examine the determinants of the cumulative survival curve S(t), and show how the shape of S(t) is directly related to the variance in the log of failure times, log(t_f_). We illustrate how changes in the variance in log(t_f_) with temperature invalidate the assumption of an average survival curve. Second, we use published data for knockdown times of adult *Drosophila* at stressful temperatures (2) to quantify how mean and variance in log of failure time vary with temperature. Our results show that for most *Drosophila* species, variance in log(t_f_) increases with increasing temperature, altering survival probabilities and the shape of S(t) between temperatures within the stressful range. Third, we propose a survival model that incorporates changes in failure variation and the shape of S(t) across temperatures, and compare the predictions with those from previous deterministic and probabilistic models in fluctuating, stressful temperature conditions using data from an experimental study with *D. melanogaster* (9). Our approach illustrates how changes in variation in failure times across stressful temperatures can alter predictions of survival in high, fluctuating temperatures. Our results highlight the importance of understanding the biological processes that contribute to variation in failure times.

## MODEL DEVELOPMENT AND RESULTS

### The survival model

The basic TDT model assumes that the mean (or median) of log(t_f_) declines linearly with increasing temperature (Fig. 1A). Variation in log(t_f_) will determine how the probability of survival changes with time. It is useful to define the **failure density f(t)**, which represents the frequency distribution of failure times for a sample of individuals. At any given temperature, the failure density and variance in log (t_f_) are related to the **cumulative survival curve S(t).** Here we use the log-logistic survival model to illustrate the main points, but the qualitative results will apply generally to other probability distributions (see Discussion). The cumulative survival function S(t) for the log-logistic model is given by:

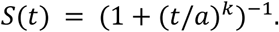

**Figure 1.**
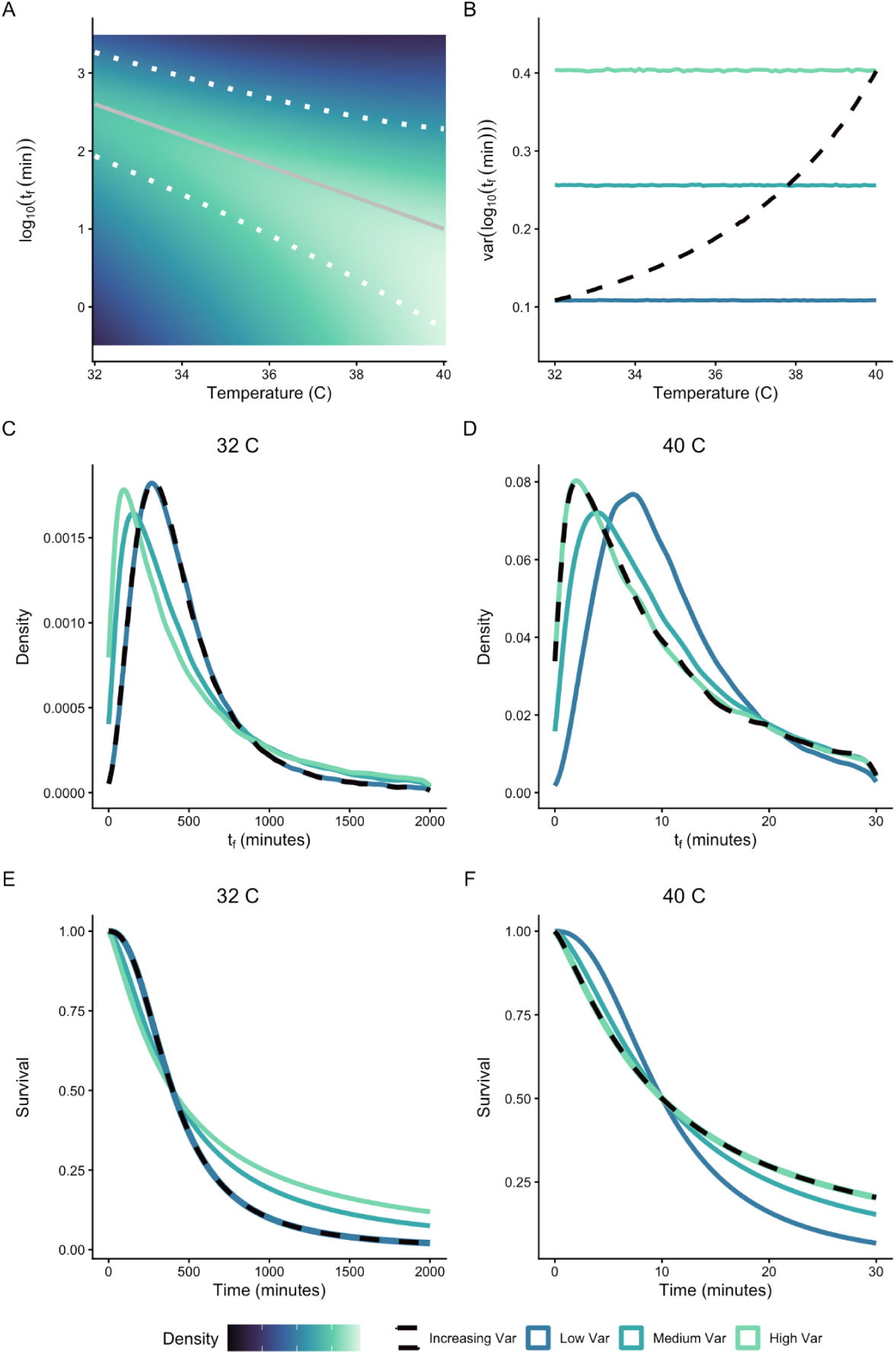
How variation in failure time t_f_ alters the failure density f(t) and cumulative survival S(t). A) Following the TDT model, let log(t_f_) decline linearly (solid gray line) with increasing temperature. Suppose that the variance in log(t_f_) increases with increasing temperature (dotted lines, 95% confidence limits; colors indicate probability densities). B) Consider four cases for variance in log(t_f_): constant variance (solid lines) at low (blue), medium (blue-green) or high (green) levels across temperature; or increasing variance with temperature (dashed black line). At any given temperature, the shape and position of the failure density (C-D) and the cumulative survival curve (E-F) varies with variance of log(t_f_).

One useful feature of the log-logistic model is that the *scale* parameter *a* equals the median of failure time t_f_. Note that the median failure time occurs when cumulative survival S(t) = 0.5 (Suppl. Fig S1). In addition, the *shape* parameter *k* is inversely related to the variance of failure time t_f_: (See Supplement):

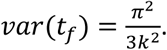

For clarity, we will italicize *scale* and *shape* when referring to the log-logistic parameters.

The **hazard function h(t)** represents the instantaneous failure rate as a function of time: i.e., the rate of failure given survival to time t (23). It is defined in terms of the failure density and the cumulative survival curve: h(t) = f(t)/S(t). In the simplest case, h(t) = λ and is constant over time, resulting in an exponential distribution. The hazard function for the log-logistic model is given by:

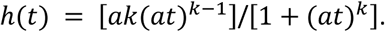

When k>1, the hazard rate increases more rapidly over time when k is larger, decreasing the width of the failure density (Suppl. Fig S1).

To explore the importance of variation in failure times, suppose the mean and median of log(t_f_) decrease linearly with increasing temperature (the TDT assumption, FIg. 1A). We will consider cases in which variance of log(t_f_) is either constant with temperature (Fig. 1B, solid lines), or increases with increasing temperature (Fig. 1A; Fig. 1B, dashed lines). Recall that variance of log(t_f_) is inversely related to the log-logistic *shape* parameter (see eqn above). The variance in log(t_f_) is therefore determined by the shapes of the failure densities and cumulative survival curves, and how these change across temperature (Fig. 1C-F; Suppl. Fig S1). At any given temperature, greater variance in log(t_f_) results in a strongly asymmetric failure density with a low mode and a long right tail (Fig. 1C-D). As a consequence, greater variance in log(t_f_) results in a more gradual decline in cumulative survival across time, whereas lower variance generates a logistic (switch-like) decline in cumulative survival (Fig. 1E-F). Because log mean failure time declines rapidly with increasing temperature (Fig. 1A), the scales of f(t) and S(t) differ strongly between temperatures. But because the variance of log(t_f_) is independent of scale (and of median log(t_f_)), any temperature effects on variance are separate from the established TDT relationship (Fig. 1C-F).

Now consider the case in which variance of log(t_f_) is not constant, but rather increases with increasing temperature (Fig. 1, dashed lines). In this case, the shapes of the failure time density and cumulative survival curve change with temperature, differing from the two constant variance cases (Fig. 1C-F). If variance of log(t_f_) changes with temperature, this has important consequences for models of failure times and survival curves in fluctuating temperature conditions (see Supplement). Deterministic TDT models such as Jørgensen *et al* (1) do not incorporate variation in failure times or the probabilistic nature of failure, and focus on predictions about median or mean failure time. Probabilistic TDT models such as Rezende *et al* (3) do not assume a specific parametric form for the cumulative survival curve S(t), but assume that there is an ‘average’ S(t) that applies across all temperatures of interest. In this sense, the model assumes that variance of log(t_f_) is constant across temperatures (see Fig. 1 and supplementary material in (3)). Below, we will address whether variance of log(t_f_) changes at different constant temperatures within the stressful range, and explore the consequences of this for predicting failure and survival in fluctuating temperatures.

### Testing TDT model assumptions: Constant temperatures

To assess how mean and variance log(t_f_) change with temperature, we used recently published datasets for knockdown times (t_f_) of adults from 11 *Drosophila* species at a series of stressful, constant temperatures (2) (Fig. 2). We excluded trials with less than 10 individuals from our analyses to ensure model convergence. We also estimated the *scale* and *shape* parameters for the log-logistic model at each temperature for each species (R package flexsurv). Preliminary analyses suggested that the log-logistic model provides a better fit for these data than alternatives such as Weibull, Log-normal, or Gompertz models.

**Figure 2.**
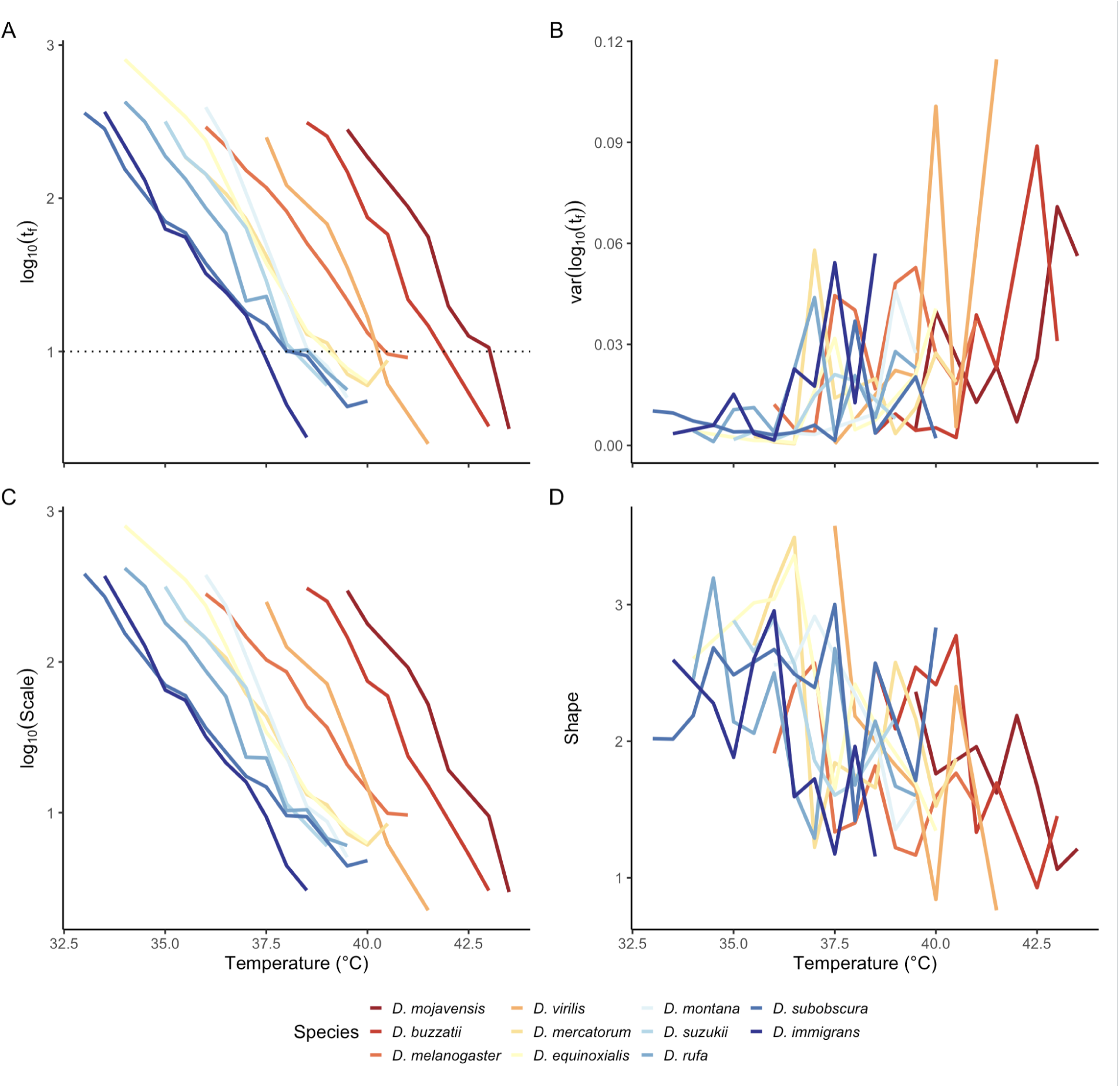
Changes in median (A) and (B) variance in adult failure (knockdown) times t_f_, and in estimated log-logistic *scale* (C) and *shape* (D) parameters, as functions of temperature for 11 *Drosophila* species. Data from (2). For each species, the median of log(t_f_) (A) and the *scale* (C) decline linearly with increasing temperature. In contrast, the variance of log(t_f_) (B) is greater and more variable at higher temperatures, and *shape* (D) generally declines with increasing temperature. Line color (blue to red) for each species is ordered using the mean of log(t_f_) at 10 minutes (dashed line in A).

As predicted by TDT models, both the mean of log(t_f_) and the value of log(*scale*) (recall that scale = median of t_f_: see previous section) decline linearly with increasing temperature (Fig. 2A, C). There are substantial differences in heat tolerance (e.g. position of x-intercept in Fig. 2A-B) among species. Residual analysis confirms that any nonlinearity in the TDT trend is small in scale and inconsistent between taxa (Suppl Fig. S2). Changes in the variance in log(t_f_) and *shape* across temperatures are more complex (Fig. 2B, D). At lower temperatures (<36℃), variance is relatively low (and *shape* is high), but the variance is generally larger at higher temperatures. Overall, *shape* declines significantly with increasing temperatures (mixed-effects LR test: *X*^2^ = 33.71, df =1, p < 0.0001), although there is variation among species in the strength of the relationship (Suppl Fig. S3). As a consequence for many species, the shape of the cumulative survival curve changes with temperature (see Supplement), contrary to the assumption of an ‘average’ survival curve across temperatures (3). We emphasize that these changes occur within a narrow range of stressfully high temperatures (>32.5℃). Interestingly, species with higher heat tolerance do not show increased variance in log(t_f_) until higher temperatures (Suppl Fig. S3), suggesting that stress responses may alter the variance and distribution of failure times.

The small sample sizes in these data (10-26 individuals per temperature) contribute to the ‘noisiness’ of our estimates of variance(log(t_f_)) and shape (Fig. 2B, D). This has important consequences for estimating failure densities and cumulative survival curves at different temperatures (Fig. 3). The Rezende *et al* (3) model estimates an average survival curve and failure density across all temperatures, and applies this at each temperature. The increasing variance log-logistic model (hereafter labeled the increasing variance model) allows the log-logistic *shape* parameter to decline linearly with temperature if such a relationship is found (Fig. 2D); otherwise *shape* is constant across temperature (eg. Fig. 3A-D). For both *D. subobscura* (Fig. 3A-D) and *D. equinoxialis* (Fig. 3 E-H), the shapes of the observed failure time densities (grey lines) differ markedly between low and high stressful temperatures and are quite jagged, reflecting the sparseness of the data. For both species, many of the observed failure times (grey lines) fall outside of the failure densities predicted by the Rezende model (orange lines), especially at the high temperature (Fig. 3A-B and 3E-F); as a consequence, the Rezende model predicts a faster decline in cumulative survival than the observed survival curve (Fig. 3C-D and 3G-H). The predicted failure densities for the increasing variance model (purple lines) are consistently wider than the observed densities, but encompass the full range of the observed failure times (Fig. 3). These patterns highlight an important difference between the models: at a given temperature, the Rezende model uses a non-parametric model fitting approach that can result in complex failure densities and survival curves, whereas the log-logistic model estimates a single parameter (*shape*) that determines the shape of the failure density and cumulative survival. The complex failure density from the Rezende model likely results from overfitting of the data, rather than reflecting the ‘true’ failure density (see Discussion).

**Fig 3.**
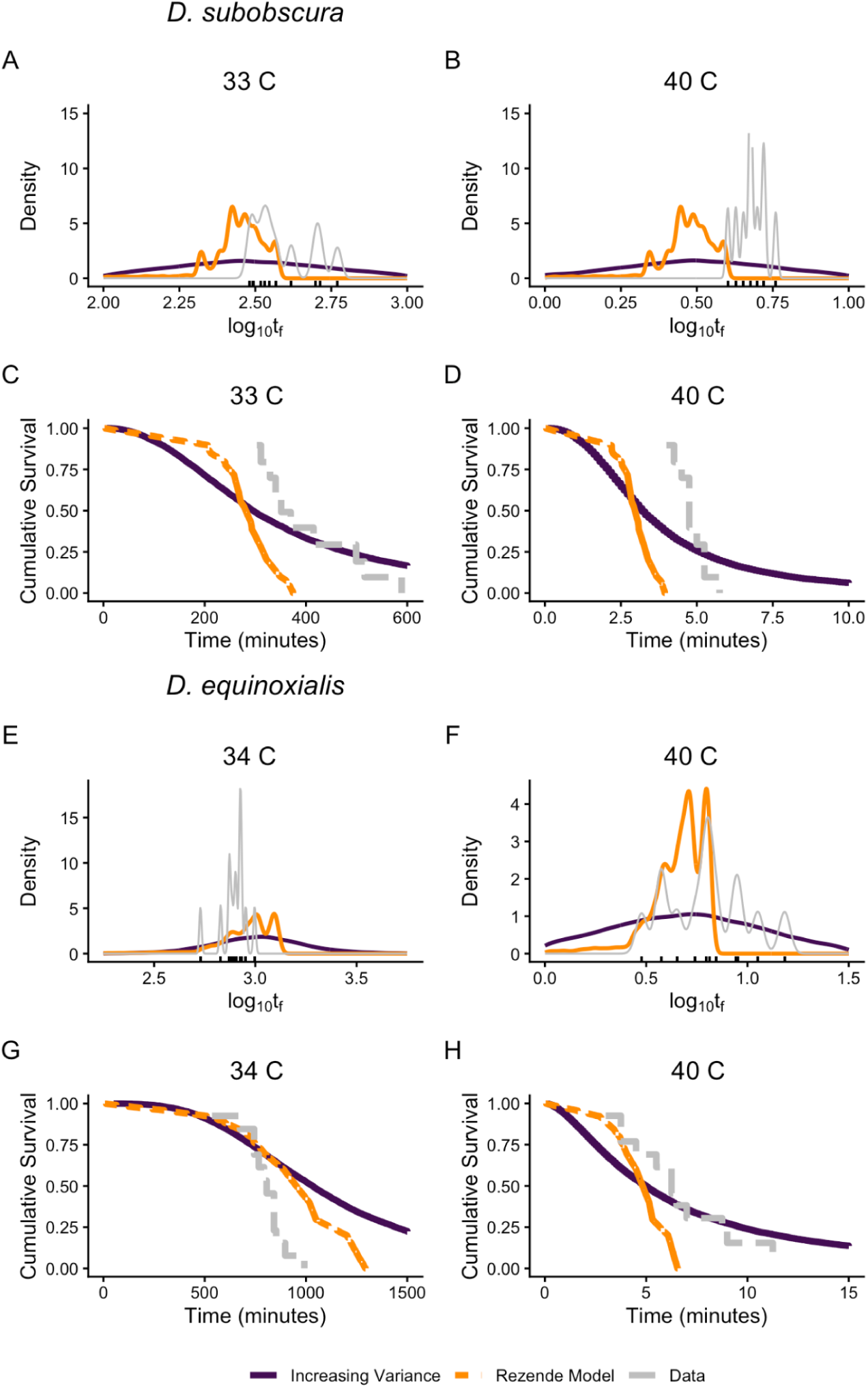
Observed and predicted failure densities (A- B) and cumulative survival curves (C-D) for *D. subobscura* (A-D) and for D. equinoxialis (E-H) at two constant temperatures. Data from (3) (hash marks in A-B and E-F). Curves for the observed data (grey lines) and predictions based on the Rezende *et al* (orange) and increasing variance (purple) models are given for each temperature. Note the greater estimated variance (purple) at the high temperature for *D. equinoxialis* (E-F) but not *D. subobscura* (A-B).

### Testing model predictions: fluctuating temperatures

How do these effects alter predicted patterns of survival in fluctuating temperature conditions? We address this question using *D. melanogaster* (a different population than that represented in Fig. 2) failure (heat coma) time data from an experiment subjecting 26 sets of adults to different fluctuating temperatures between 34 and 42℃ (1). Jørgensen *et al* (1) used their deterministic TDT model, assuming that injury accumulation is additive, to predict median failure time (t_f_) for each set of adults; we compared their predictions along with predictions for the Rezende *et al* and the increasing variance (log-logistic) models to the observed failure times (Fig. 4). Based on the sum of the (absolute) errors the increasing variance (187.2) and Jørgensen (188.8) models yield better predictions for median failure time of males than the Rezende model (222.5). For females, the Jørgensen model (189.7) has lower error than the increasing variance model (194.7), and both have much lower error than the Rezende model (276.7). Note that at lower mean temperatures (e.g. t_f_ > 150 minutes) all three models consistently underestimate the observed median t_f_ (Fig. 4). For the *D. melanogaster* population used in this experiment, the relationship between temperature and shape was weak, so we also compared log-logistic models with constant or with increasing variance. Incorporating increasing variance had little effect on the predictions for the log-logistic model (Fig. S5).

**Fig. 4.**
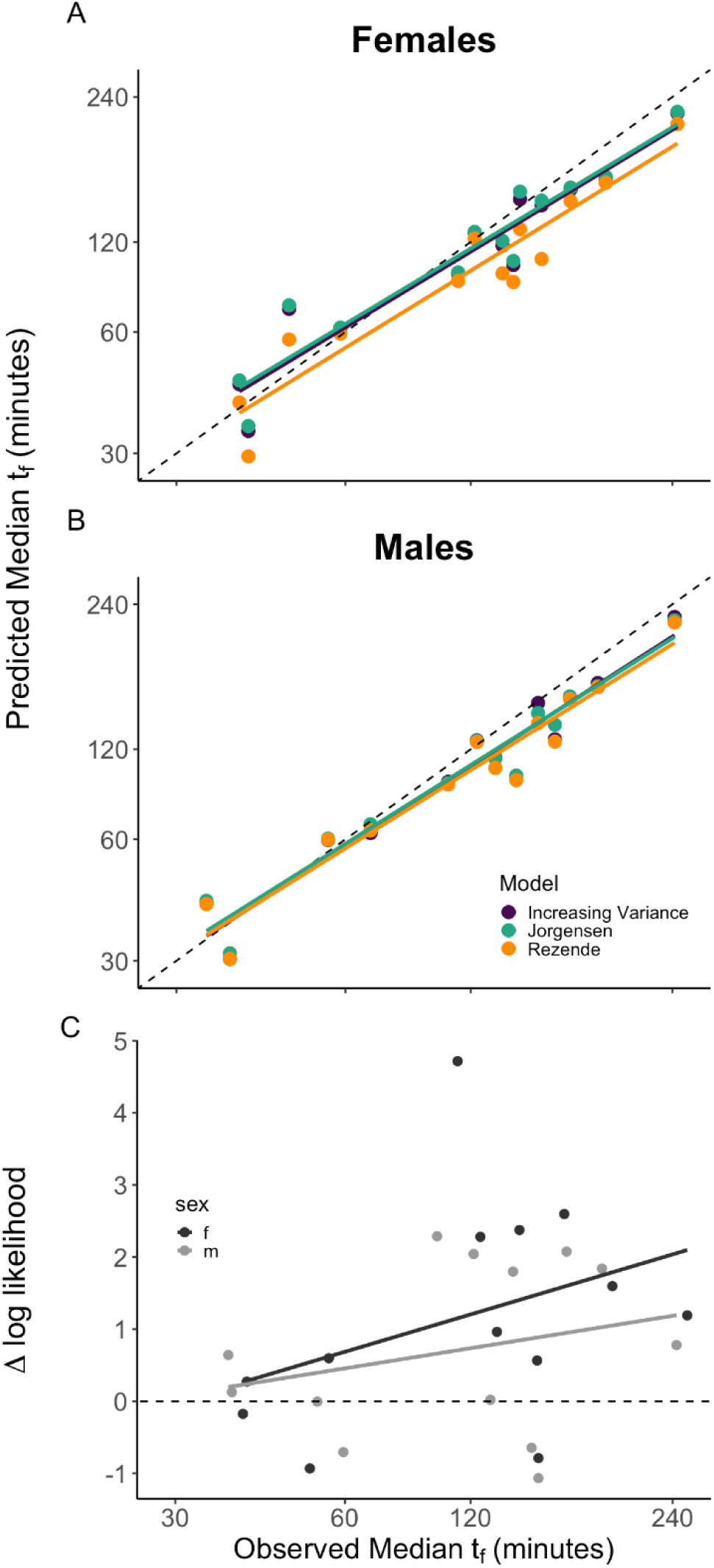
Observed vs predicted median failure times (A-B) and survival curves (C) for sets of adult *D. melanogaster* in different fluctuating stressful temperature conditions. Data from (1). Panel A: Females; Panel B: Males. Predictions for the Jørgensen *et al* (teal), Rezende *et al* (orange) and log-logistic increasing variance (purple) models. Linear regression line for each model is shown, as well as the 1:1 line (light grey line). Panel C: Difference in log-likelihood of the increasing variance relative to the Rezende model for predicting the cumulative survival curves, plotted as a function of observed median failure time. Positive values indicate a better fit for the increasing variance model. Values for females (purple) and males (teal) are given separately; regression lines (non-significant) are also indicated.

The probabilistic models can also predict the distribution of failure times (and hence the cumulative survival curves) under fluctuating temperatures for each set of individuals. The difference in log-likelihood between the increasing variance (log-logistic) and Rezende models quantified the relative fit of the two models for each set (Fig. 4C) (see Supplement for details). Based on log-likelihood values computed for each set, the increasing variance (log-logistic) model provides a significantly better fit than the Rezende model (paired t-test: t = 3.4685, df = 25, p = 0.0019), with a higher likelihood in 20 of 26 cases. Together these results suggest that the (deterministic) Jørgensen model and the log-logistic model yield better predictions of median failure times than the Rezende model, and the log-logistic model yields better predictions of variation in failure times and in cumulative survival than the Rezende model.

## DISCUSSION

### Modeling survival at stressful temperatures

The thermal death time (TDT) model- specifically, that the average time to death or failure declines exponentially with increasing temperature– was first proposed for microbes more than a century ago (13), and subsequently confirmed for fish and insects (14,21) more than a half-century ago. Numerous recent studies have documented the TDT relationship in a variety of ectothermic organisms at constant, stressful temperatures (2,15,16,18–20,26–28). Recent extensions enable applying the TDT model to fluctuating temperatures but require additional assumptions about the joint effects of time and temperature on survival probabilities or failure rates. Deterministic models assume that failure at stressful temperatures occurs when damage accumulation exceeds some critical threshold, and that damage accumulation at different (stressful) temperatures is additive (1,2,9); experimental evidence in several systems supports the additivity of damage accumulation (1,9,14,28). These deterministic models can be used to predict average failure times in fluctuating temperatures– an important contribution–but ignore the stochastic nature of failure and cannot directly predict survival probabilities or survival curves. Alternatively, the influential probabilistic model of Rezende et al (3) uses failure data at different constant temperatures to compute an average cumulative survival curve, implicitly assuming that the shape of the survival curve, and variation in failure time, does not change across temperatures. Here we have explored these two assumptions and their consequences for modeling survival and failure in both constant and fluctuating temperatures. We show that median log(t_f_) (and log-logistic *scale*) decline linearly with increasing temperature, confirming previous analyses in support of the TDT model. In contrast, for most species, var[log(t_f_)] and *shape* change in magnitude and variability across temperatures (Fig. 2). In particular, var[log(t_f_)] significantly increases (shape significantly decreases) with increasing temperatures for many species, even over the narrow range of stressful temperatures considered in these studies (Fig. 2). Increasing var[log(t_f_)] with temperature (Fig. 2) will alter the shape and curvature of the cumulative survival curve (Fig. 1 and S1), contrary to the constant shape assumption (3). Note however that the presence or strength of the relationship between temperature and var[log(t_f_)] differs substantially among the *Drosophila* species and populations considered here (Fig. S3). The estimated values of the log-logistic *shape* parameter for *Drosophila* at high, fluctuating temperatures–typically in the range of 1-3 (Fig. 2)-- predict hazard functions that either decline or increase gradually with time (Fig. S1), contrary to expectations for a deterministic threshold model of failure.

We have used the log-logistic survival model to illustrate our points, but the main findings will apply to other probability distributions. One benefit of the log-logistic model is that its parameters– *scale* and *shape*– are readily related to the mean and variation in failure times t_f_ : *scale* equals the median of log(t_f_), and *shape* declines with increasing variance in log(t_f_). A key insight is that var[log(t_f_)] determines the shape of the survival curve: cumulative survival declines more gradually as the variance in log(t_f_) increases (Fig. 1 and S1). These considerations highlight the importance of understanding both the mean and variance in failure times for modeling survival.

### Understanding variation in failure times

Our estimates of var[log(t_f_)] and *shape* here are quite imprecise and noisy (Fig. 2), as a consequence of the small sample sizes in these data (10-26 individuals per temperature). For data sets with small sample sizes, the empirically-based Rezende *et al*. approach can result in predicted failure densities and survival curves that fail to overlap much of the observed data at particular temperatures (Fig. 3). Substantially larger sample sizes will be needed to provide better estimates of variance in log(t_f_) and of *shape* parameters, and to select the best parametric models for failure time and survival data.

Why does var[log(t_f_)] change with temperatures within the stressful range? One possibility is that the temporal effects of stress on failure (knockdown or mortality) may vary within the stressful temperature range. For example, if different mechanisms of stress response, such as HSP production, ROS accumulation, and DNA repair are induced at different stressful temperatures, this could generate differences in failure densities across temperatures (11,29,30). Variation among individuals in their responses to stressful temperatures could also generate this pattern. The probabilistic models considered here assume random variation among individuals across all temperatures: in particular, by assuming constant variance across temperatures, TDT models imply that individuals vary in the intercept but not the slope of the TDT relationship. Alternatively, variation among individuals or genotypes in the slope (more generally shape) of their TDT curves can generate changes in var[log(t_f_)], and therefore *shape*, with temperature. While some studies have examined genetic variation in thermal tolerance (31), phenotypic and genetic variation in TDT curves is largely unexplored (but see (32,33)); studies with clones or isogenic lines may be particularly valuable to addressing this issue.

Studies to date have emphasized estimates or predictions of mean or median failure times. However, many important biological questions require information about cumulative survival curves (equivalently, variation in failure times). Calculations of population growth rate (e.g. r) require information about the survival curves, as do predictions about population persistence or extinction (34,35). Similarly, the shape of survival curves is important for predicting the survival of immature states to adulthood, especially when environmental factors (including temperature) influence both developmental and mortality rates. It is thus important for analyses and models to move beyond the current focus on average TDTs.

### Modeling survival in fluctuating temperatures

Using constant temperature studies to estimate the parameters for models that can predict survival in fluctuating temperatures is a powerful approach, and such predictions can be experimentally tested in controlled environmental conditions (1,2,14). We find that the Jørgensen and increasing variance (log-logistic) models yield substantially better predictions for median failure time than the Rezende model (Fig. 4). In addition, the model comparison shows that the increasing variance (log-logistic) model yields a significantly better fit to the observed failure densities than the Rezende model (Fig. 4). Increased variance made only minor contributions to the superiority of the log-logistic model for these data (Fig. S5). The weaker performance of the Rezende model here may be a consequence of overfitting of the average failure density and cumulative survival curve (Fig. 3) based on limited data, resulting in less accurate predictions. While larger sample sizes would help resolve this, further discussion of the relative merits of parametric and non-parametric survival models seems warranted here.

Importantly, all three fluctuating temperature models consistently underestimate median failure times when the frequency of lower (but still stressful) temperatures is greater– i.e. at longer median failure times (Fig. 4). Residual analysis (Suppl Fig. S1) supports the linear relationship between mean[log(t_f_)] and temperature over this temperature range, as assumed by TDT models. Instead, the assumption that damage accumulation is additive may not apply when temperatures vary across the stressful range.

Studies exposing fish, insects, and plants to various pairs of two successive stressful temperatures suggest that median t_f_ can be accurately predicted with the additive stress model (1,14,19,28). However, these experiments only tested by switching between pairs of temperatures, have spanned only a narrow portion of the stressful range, and did not consider variance in failure times. For example, in one such study with *D. melanogaster,* variance in failure time is much higher when the lower temperature (36.5℃) preceded the higher (39.5℃), compared with the reverse order (1). Such a result indicates that the order of temperature exposure affects failure times, and may result from acclimation, recovery or repair (1,11,12,36). Additional tests of additivity will improve our understanding of these processes and inform further development of TDT models.

### Beyond stressful temperatures

The models and data described here consider a restricted range of stressfully high temperatures, in which failure typically occurs within minutes to a few hours (Figs. 2-4). Of course, most terrestrial thermal environments include diurnal and other fluctuations between stressful and non-stressful (‘permissive’) temperatures. A growing literature explores whether and how exposure to non-stressful temperatures allows repair and recovery of damage accumulated during thermal stress (9,12,29,37). Some recent experimental studies show that cool night-time temperatures can reduce damage accumulated at stressful high daytime temperatures, and improve survival and growth (38–40). Incorporating the time-dependent effects of recovery and repair into models of responses to diurnally fluctuating temperatures represents an important challenge (41,42). Multiple deterministic approaches have been proposed for modeling the time-dependent effects of damage accumulation, recovery, and repair to predict mean survival or growth in fluctuating conditions (9,19,36,37,41,42).

Accurately predicting survival in variable field conditions poses additional challenges (3,37,43), although initial attempts are encouraging (3). Recent syntheses have shown that the TDT slope is steeper within the stressful range than within the permissive range for many ectotherms (9,15), but these analyses have considered only averages and not variation in failure times. Survival curves and failure densities at non-stressful temperatures will be determined by aging, ontogeny, acclimation and other processes, rather than damage accumulation (21,22). One approach is to assume that failure rates at permissive temperatures are constant over time, implicitly assuming that survival is exponentially distributed at these temperatures. Because knockdown and related assays only apply at stressful temperatures, death or other metrics of failure that apply across the full range of temperatures may be more useful in this regard (14,21). A related issue is how to identify the threshold temperature that distinguishes permissive and stressful temperature ranges, and whether these represent distinct physiological ranges rather than part of a continuum (15,36). Given these many challenges, experimental tests with fluctuating temperatures in environmental chambers may be a particularly useful approach to evaluating alternative models for predicting survival in variable environments (9,36).

## ACKNOWLEDGEMENTS

We thank Lorrie He, Katherine Malinski, Tyler Pereira, Chris Willett, and two anonymous reviewers for feedback on previous drafts of the manuscript; Geoff Legault and Mete Yuksel for statistical advice; Michael Orsted and Johannes Overgaard for access to data; and the Whiteley Center at Friday Harbor Labs (University of Washington) for providing an ideal venue for working on this project.

## AUTHOR CONTRIBUTIONS

Conceptualization: JGK, LBB. Data curation: LBB, GWB. Formal analysis: GWB, JGK. Funding acquisition: JGK. Investigation: all. Methodology: all. Software: GWB. Supervision: JGK. Visualization: all. Writing – original draft: JGK, LBB. Writing – review & editing: all.

## METHODS AND DATA AVAILABILITY

Code and Data used in the figures and analyses are available at: https://github.com/gbullard87/Beyond_Mean_TDT_2025.

## ACKNOWLEDGEMENTS

We thank Lorrie He, Katherine Malinski, Tyler Pereira, Chris Willett, and two anonymous reviewers for feedback on previous drafts of the manuscript; Michael Ørsted and Johannes Overgaard for access to data; and the Whiteley Center at Friday Harbor Labs (University of Washington) for providing an ideal venue for working on this project.

## REFERENCES

1. Jørgensen LB, Malte H, Ørsted M, Klahn NA, Overgaard J. A unifying model to estimate thermal tolerance limits in ectotherms across static, dynamic and fluctuating exposures to thermal stress. Sci Rep. 2021 June 18;11(1):12840.

2. Jørgensen LB, Malte H, Overgaard J. How to assess *Drosophila* heat tolerance: Unifying static and dynamic tolerance assays to predict heat distribution limits. White C, editor. Functional Ecology. 2019 Apr;33(4):629–42.

3. Rezende EL, Bozinovic F, Szilágyi A, Santos M. Predicting temperature mortality and selection in natural Drosophila populations. Science. 2020 Sept 4;369(6508):1242–5.

4. Bennett JM, Calosi P, Clusella-Trullas S, Martínez B, Sunday J, Algar AC, et al. GlobTherm, a global database on thermal tolerances for aquatic and terrestrial organisms. Sci Data. 2018 Mar 13;5(1):180022.

5. Sunday JM, Bates AE, Kearney MR, Colwell RK, Dulvy NK, Longino JT, et al. Thermal-safety margins and the necessity of thermoregulatory behavior across latitude and elevation. Proceedings of the National Academy of Sciences. 2014 Apr 15;111(15):5610–5.

6. Clusella-Trullas S, Garcia RA, Terblanche JS, Hoffmann AA. How useful are thermal vulnerability indices? Trends in Ecology & Evolution. 2021 Nov 1;36(11):1000–10.

7. Terblanche JS, Deere JA, Clusella-Trullas S, Janion C, Chown SL. Critical thermal limits depend on methodological context. Proc Biol Sci. 2007 Dec 7;274(1628):2935–43.

8. Kingsolver JG, Umbanhowar J. The analysis and interpretation of critical temperatures. Journal of Experimental Biology. 2018 June 27;221(12):jeb167858.

9. Ørsted M, Jørgensen LB, Overgaard J. Finding the right thermal limit: a framework to reconcile ecological, physiological and methodological aspects of CTmax in ectotherms. Journal of Experimental Biology. 2022 Oct 3;225(19):jeb244514.

10. Rezende EL, Castañeda LE, Santos M. Tolerance landscapes in thermal ecology. Fox C, editor. Functional Ecology. 2014 Aug;28(4):799–809.

11. González-Tokman D, Córdoba-Aguilar A, Dáttilo W, Lira-Noriega A, Sánchez-Guillén RA, Villalobos F. Insect responses to heat: physiological mechanisms, evolution and ecological implications in a warming world. Biological Reviews. 2020;95(3):802–21.

12. Williams CM, Buckley LB, Sheldon KS, Vickers M, Pörtner HO, Dowd WW, et al. Biological Impacts of Thermal Extremes: Mechanisms and Costs of Functional Responses Matter. Integrative and Comparative Biology. 2016 July 1;56(1):73–84.

13. Bigelow WD. The logarithmic nature of thermal death time curves. The Journal of Infectious Diseases. 1921 Nov 1;29(5):528–36.

14. Fry, F. E. J., Hart, J. S., Walker, K. F. Lethal temperature relations for a sample of young speckled trout, Salvelinus fontinalis. Publications of the Ontario Fisheries Research Laboratory. 1946;66:9–35.

15. Jørgensen LB, Ørsted M, Malte H, Wang T, Overgaard J. Extreme escalation of heat failure rates in ectotherms with global warming. Nature. 2022 Nov;611(7934):93–8.

16. Tarapacki P, Jørgensen LB, Sørensen JG, Andersen MK, Colinet H, Overgaard J. Acclimation, duration and intensity of cold exposure determine the rate of cold stress accumulation and mortality in *Drosophila suzukii*. Journal of Insect Physiology. 2021 Nov 1;135:104323.

17. Holmstrup M, Touzot M, Slotsbo S. Characterization of the thermal death time landscape for *Enchytraeus albidus*. Pedobiologia. 2023 June 1;97–98:150876.

18. Li YJ, Chen SY, Jørgensen LB, Overgaard J, Renault D, Colinet H, et al. Interspecific differences in thermal tolerance landscape explain aphid community abundance under climate change. Journal of Thermal Biology. 2023 May 1;114:103583.

19. Ørsted M, Willot Q, Olsen AK, Kongsgaard V, Overgaard J. Thermal limits of survival and reproduction depend on stress duration: A case study of Drosophila suzukii. Ecology Letters. 2024;27(3):e14421.

20. Wehrli M, Slotsbo S, Ge J, Holmstrup M. Acclimation temperature influences the thermal sensitivity of injury accumulation in *Folsomia candida* at extreme low and high temperatures. Current Research in Insect Science. 2024 Jan 1;6:100089.

21. Hollingsworth MJ. Temperature and length of life in *Drosophila*. Experimental Gerontology. 1969 Mar 1;4(1):49–55.

22. Ricklefs RE, Scheuerlein A. Biological Implications of the Weibull and Gompertz Models of Aging. The Journals of Gerontology: Series A. 2002 Feb 1;57(2):B69–76.

23. Ergon T, Borgan Ø, Nater CR, Vindenes Y. The utility of mortality hazard rates in population analyses. Methods in Ecology and Evolution. 2018;9(10):2046–56.

24. Wilson DL. The analysis of survival (mortality) data: Fitting Gompertz, Weibull, and logistic functions. Mechanisms of Ageing and Development. 1994 May 1;74(1):15–33.

25. Cook AM, Rezende EL, Petrou K, Leigh A. Beyond a single temperature threshold: Applying a cumulative thermal stress framework to plant heat tolerance. Ecology Letters. 2024;27(3):e14416.

26. Peralta-Maraver I, Rezende EL. Heat tolerance in ectotherms scales predictably with body size. Nat Clim Chang. 2021 Jan;11(1):58–63.

27. Villeneuve AR, White ER. Predicting organismal response to marine heatwaves using dynamic thermal tolerance landscape models. Journal of Animal Ecology [Internet]. 2024 [cited 2025 June 19];n/a(n/a). Available from: https://onlinelibrary.wiley.com/doi/abs/10.1111/1365-2656.14120

28. Faber AH, Ørsted M, Ehlers BK. Application of the thermal death time model in predicting thermal damage accumulation in plants. Journal of Experimental Botany. 2024 June 7;75(11):3467–82.

29. Colinet H, Sinclair BJ, Vernon P, Renault D. Insects in Fluctuating Thermal Environments. Annual Review of Entomology. 2015 Jan 7;60(Volume 60, 2015):123–40.

30. Rennolds CW, Bely AE. Integrative biology of injury in animals. Biological Reviews. 2023;98(1):34–62.

31. Logan ML, Cox RM, Calsbeek R. Natural selection on thermal performance in a novel thermal environment. Proceedings of the National Academy of Sciences. 2014 Sept 30;111(39):14165–9.

32. Leiva FP, Santos M, Niklitschek, Edwin J., Rezende EL, Wilco C. E. P. Verberk. Genetic variation of heat tolerance in a model ectotherm: an approach using thermal death time curves. ecoevorxiv.org; 2025.

33. Soto J, Pinilla F, Olguín P, Castañeda LE. Genetic Architecture of the Thermal Tolerance Landscape in Drosophila melanogaster. Molecular Ecology. 2025;34(7):e17697.

34. Beissinger SR, McCullough DR, editors. Population Viability Analysis [Internet]. Chicago, IL: University of Chicago Press; 2002 [cited 2025 June 19]. 593 p. Available from: https://press.uchicago.edu/ucp/books/book/chicago/P/bo3637258.html

35. Roff D. Life History Evolution. Oxford, New York: Oxford University Press; 2001. 527 p.

36. Buckley LB, Huey RB, Ma CS. How damage, recovery, and repair alter the fitness impacts of thermal stress. Integrative and Comparative Biology. 2025 May 13;icaf019.

37. Arnold PA, Noble DWA, Nicotra AB, Kearney MR, Rezende EL, Andrew SC, et al. A Framework for Modelling Thermal Load Sensitivity Across Life. Global Change Biology. 2025;31(7):e70315.

38. Ma CS, Wang L, Zhang W, Rudolf VHW. Resolving biological impacts of multiple heat waves: interaction of hot and recovery days. Oikos. 2018;127(4):622–33.

39. Zhao F, Zhang W, Hoffmann AA, Ma CS. Night warming on hot days produces novel impacts on development, survival and reproduction in a small arthropod. Journal of Animal Ecology. 2014;83(4):769–78.

40. Bai CM, Ma G, Cai WZ, Ma CS. Independent and combined effects of daytime heat stress and night-time recovery determine thermal performance. Biology Open. 2019 Mar 26;8(3):bio038141.

41. Klanjscek T, Muller EB, Nisbet RM. Feedbacks and tipping points in organismal response to oxidative stress. Journal of Theoretical Biology. 2016 Sept 7;404:361–74.

42. Kingsolver JG, Woods HA. Beyond Thermal Performance Curves: Modeling Time-Dependent Effects of Thermal Stress on Ectotherm Growth Rates. The American Naturalist. 2016 Mar;187(3):283–94.

43. Buckley LB, Kingsolver JG. Evolution of Thermal Sensitivity in Changing and Variable Climates. Annual Review of Ecology, Evolution, and Systematics. 2021 Nov 3;52(Volume 52, 2021):563–86.

